# Methods for chair restraint and training of the common marmoset on oculomotor tasks

**DOI:** 10.1101/230235

**Authors:** Kevin D. Johnston, Kevin Barker, Lauren Schaeffer, David Schaeffer, Stefan Everling

**Affiliations:** Department of Physiology and Pharmacology, Schulich School of Medicine and Dentistry, University of Western Ontario, London, Ontario, Canada; Brain and Mind Institute, University of Western Ontario, London, Ontario, Canada; Neuronitek, London, Ontario, Canada; Robarts Research Institute, London, Ontario, Canada

**Keywords:** common marmoset, saccade, oculomotor, gap effect, neurophysiology

## Abstract

The oculomotor system is the most thoroughly understood sensorimotor system in the brain, due in large part to electrophysiological studies carried out in macaque monkeys trained to perform ocuolomotor tasks. A disadvantage of the macaque model is that many cortical oculomotor areas of interest lie within sulci, making high-density array and laminar recordings impractical. Further, many techniques of molecular biology developed in rodents, such as transgenic animals and optogenetic manipulation of neuronal subtypes, are limited in this species. The common marmoset (*Callithrix jacchus*) may potentially bridge the gap between systems neuroscience in macaques and molecular techniques, and additionally possesses a smooth cortex allowing easy access to frontoparietal oculomotor areas. To date, techniques for restraint and training of these animals to perform oculomotor tasks remain in an early stage of development. Here we provide details of a custom-designed chair for restraint of marmosets, a combination head restraint/recording chamber providing stability suitable for eye movement and neural recordings, and a training protocol for oculomotor tasks. As proof-of-principle, we report the results of a psychophysical study in marmosets trained to perform a saccade task using these methods, showing that, as in rhesus and humans, marmosets exhibit a “gap effect” – a decrease in reaction time when the fixation stimulus is removed prior to the onset of a visual saccade target. These results provide evidence that the common marmoset is a suitable model for neurophysiogical investigations of oculomotor control.

**NEW AND NOTEWORTHY:** The ability to carry out neuronal recordings in behaving primates has provided a wealth of information regarding the neural circuits underlying the control of eye movements. Such studies require restraint of the animal within a primate chair, head fixation, methods of acclimating the animals to this restraint, and the use of operant conditioning methods for training on oculomotor tasks. In contrast to the macaque model, relatively few studies have reported in detail methods for use in the common marmoset. Here we detail custom-designed equipment and methods by which we have used to successfully train head-restrained marmosets to perform basic oculomotor tasks.

## INTRODUCTION

The common marmoset (*Callithrix jacchus*) is a small-bodied New World primate that is rapidly gaining popularity as a model animal for biomedical research (Mansfield, 2003). More specifically, this species possesses numerous advantages as a model for neuroscience research. Marmosets share a homologous network of cortical and subcortical structures with humans (Solomon & Rosa, 2014; Bakola et al., 2015; Burkart & Finkenwirth, 2015; Ghahremani et al., 2017). Transgenic (Sasaki et al., 2009) and gene-editing (Kishi et al., 2014) tools have been successfully developed in this species (see for review Sasaki et al., 2015), as have optogenetic approaches to manipulating specific elements of neural circuits (MacDougall et al, 2016). These animals can also be operantly conditioned to perform cognitive tasks designed to probe abilities such as set-shifting (Dias et al, 1996; Dias et al, 1997; Roberts & Wallis, 2000) and working memory (Collins et al., 1998; Spinelli et al., 2004; Takemoto et al., 2011; Yamazaki et al., 2016), thus allowing the possibility of combining genetic and behavioural neuroscience approaches in a primate species. Large-scale brain mapping projects to elucidate the detailed anatomy of the marmoset brain, such as the Brain/MINDS Japanese national brain project are in progress (Okano & Mitra, 2015; Okano et al., 2016). Finally, the marmoset cortex is relatively lissencephalic, which offers practical advantages for neurophysiological investigations. Areas of interest are easily accessible on the dorsal cortical surface, making it possible to carry out high-density electrophysiological recordings using planar or laminar electrode arrays (Mitchell & Leopold, 2015). This property of the marmoset brain has also been exploited extensively for single electrode studies investigating auditory processing using the marmoset model (Lu et al., 2001; Barbour & Wang, 2003; Wang et al., 2005; Bendor & Wang, 2005; Bendor & Wang, 2007).

The oculomotor network is perhaps the most thoroughly understood sensorimotor system in the brain, owing largely to the wealth of neurophysiological studies carried out in Old World macaque monkeys (Luschei & Fuchs, 1970; Wurtz & Goldberg, 1972; Hikosaka et al., 1983; Bruce & Goldberg, 1985; Schlag & Schlag-Rey, 1987; Andersen et al., 1987; Funahashi et al., 1989). Adapting the marmoset model to the study of oculomotor control presents some unique advantages. Cortical areas of the oculomotor circuit inaccessible to laminar recordings in the macaque due to their location within sulci, such as the frontal eye fields (FEF), are accessible in the marmoset, allowing the study of laminar circuits. This is critical as the majority of cortical synapses lie within rather than across cortical columns (Binzegger et al., 2004). Studies of oculomotor control also have a long history within neuropsychiatric research (Diefendorf & Dodge, 1908; see for review Klein & Ettinger, 2008), with many patient populations exhibiting characteristic performance deficits in tasks such as antisaccades (Fukushima et. al, 1988), and memory-guided saccades (Park & Holzman, 1992; see for review Gooding & Basso, (2008). The potential of carrying out combined behavioural and neurophysiological investigations together with genetic manipulations holds substantial promise for extending our knowledge of neural circuit changes in neuropsychiatric disorders.

The marmoset can be considered a suitable model for studies of oculomotor control on the basis of structural and functional similarity to humans and macaques (Preuss, 2000). Marmosets are foveate animals with a well-developed visual system (Solomon & Rosa, 2014; Mitchell & Leopold, 2015), and both anatomical (Collins et al.2005) and resting-state fMRI (Ghahremani et al., 2017) evidence has established homologous frontal and parietal areas with direct connections to the midbrain superior colliculus (SC), an oculomotor structure critical for saccade initiation. Saccades can be evoked by microstimulation within a subregion of frontal cortex in this species (Mott et al., 1910; Blum et al., 1982; Burish et al., 2008), further suggesting the existence of a FEF homologous to that of macaques (Bruce & Goldberg, 1985) and humans (Paus, 1996). Functionally, marmosets can be trained to make saccades (Mitchell et al., 2014; Nummela et al., 2016) and smooth pursuit eye movements (Mitchell et al., 2015) with metrics comparable to macaques and human participants, though they exhibit a decreased oculomotor range (Mitchell et al., 2014).

Oculomotor research using the behaving macaque, and the procedures for carrying out this research, have developed over a period of close to fifty years (Wurtz, 1969). In contrast, neurophysiological studies in behaving marmosets have a much shorter history. Procedures for chair restraint for the purposes of behavioural study were first developed in the late 1980’s (O’Byrne & Morris, 1988), however head restraint and simultaneous recordings in conscious animals have only been carried out much more recently (Lu et al., 2001), and these procedures combined with a performance of a behavioural task, more recently still (Remington et al., 2012). Head restraint and combined oculomotor training have only been conducted within the last five years (Mitchell et al., 2014). Accordingly, the extant literature on the design of primate chairs, systems for head restraint, and training procedures, both for behavioural acclimation to chair restraint, and on oculomotor tasks, is much less extensive, which presents a challenge to researchers wishing to adopt this model. Moreover, the much smaller physical size of the marmoset presents some practical difficulties with respect to directly adapting techniques developed in the macaque model. For example, implantation of recording chambers to access brain areas of interest in addition to head restraint bolts can be challenging due to the small size of the head.

Here, we provide details of an integrated custom-designed marmoset chair, and implantable combination head restraint device / recording chamber we have used in the successful training of marmosets, and for neurophysiological recordings during oculomotor tasks in the behaving marmoset. We additionally describe our procedures for acclimation to chair restraint, and training on basic saccade tasks. Finally, we present behavioural data from animals trained using this system, replicating in the marmoset the gap effect (Saslow, 1967), an oculomotor phenomenon well established in both humans (Fischer & Ramsperger, 1984) and macaques (Fischer & Boch, 1983; Munoz et al., 2000).

## MATERIALS AND METHODS

### Subjects

All training and surgical procedures described below were carried out on two male common marmosets (*Callithrix jacchus),* weighing 340gm (Marmoset M) and 512gm (Marmoset B), and aged 4 and 2 years, respectively, at the time of the experiments. All experimental methods described were performed in accordance with the guidelines of the Canadian Council on Animal Care policy on the care and use of experimental animals and an ethics protocol approved by the Animal Care Committee of the University of Western Ontario.

### Combination head restraint / recording chamber

Conducting electrophysiological recordings in combination with oculomotor behaviour requires a method for maintaining fixation of the head, and a compatible method allowing chronic access to the brain. Stability of the system is critical to ensure the stability of both behavioural and neuronal recordings. In the macaque model, this has been accomplished by constructing an extracranial implant consisting of a plastic or metal head restraint post or bolt, and recording chambers fixed in place by skull screws and methyl methacrylate dental resin (Komatsu & Wurtz, 1988; DeSouza & Everling, 2004; Overton et al., 2017). The head post is attached to the primate chair to ensure stabilization, and single or multiple electrode microdrives to the recording chambers. This method of recording is not practical in marmosets due to their small size. Many commercially available microdrive systems are designed to fit recording chambers of appropriate size for macaques, which are too large for use in the marmoset, and the weight of these systems is substantial. The use of implanted planar electrode arrays such as the Utah array has been carried out in the marmoset (Chaplin et al., 2017), but the fixed nature of the array does not allow movement of electrodes in order to better isolate the extracellular potentials of single neurons. Acrylic implants with head posts similar to those employed in macaques have been employed in studies of marmoset auditory cortex, with access to the brain provided by burr holes opened above the cortex and surrounded by a thin wall of acrylic, with the microelectrode drive mounted separately from the animal’s head (Lu et al.2001). Behavioural studies of oculomotor control in the marmoset have used a similar post system for head restraint (Mitchell et al., 2014; Mitchell et al., 2015; Nummela et al., 2017).

A primary interest of our laboratory is the investigation of the role of frontoparietal circuits in oculomotor control (Johnston and Everling, 2008). To electrophysiologically interrogate such circuits in the marmoset at the level of single neurons, we reasoned that simultaneous recordings of frontal and parietal cortex with moveable electrodes would be advantageous. This requires access to the brain at up to four locations simultaneously, across a large area of the marmoset skull. For such studies, a head restraint system consisting of a headpost presents some disadvantages. First, the post requires stable attachment to the head, necessitating a large acrylic “footprint” potentially rendering access to areas of interest difficult. Second, due to the small size of the marmoset head and the logistics of positioning multiple electrodes within micromanipulators at different sites across the skull, a headpost could prevent a physical barrier rendering multiple simultaneous recordings impossible. To avoid these issues, we adapted a system used in early neurophysiological investigations in the rhesus model (Porter et al., 1971; Lemon, 1984), which surrounds the skull in a halo or ring-like fashion allowing both head restraint and access to the skull inside the ring. We constructed from polyether ether ketone (PEEK) plastic an oval-shaped recording chamber of sufficient length and width to cover and allow access to a substantial anteroposterior and medio-lateral extent of the marmoset skull. The base of the chamber is curved to match the contour of a model skull to allow for a better fit. A schematic of this chamber is presented in Figure 1A, and marmoset skull fitted with this system is presented in Figure 1B. Within this design, we incorporated four conical receptacles, placed laterally on both sides at the anterior and posterior extent of the chamber. These were designed to mate with the male ends of four moveable fixating pins attached to brackets mounted on a custom-built stereotaxic frame. The frame allows precise mounting of multiple micromanipulators for advancing electrodes, and the common mounting points of the skull with chamber and micromanipulators provide recording stability. Burr holes drilled within the chamber during a surgical procedure allow access to multiple cortical areas simultaneously. The chamber is covered with a custom designed cap fixed in place with set screws. Surgical details are presented in detail below.

**Figure 1.**
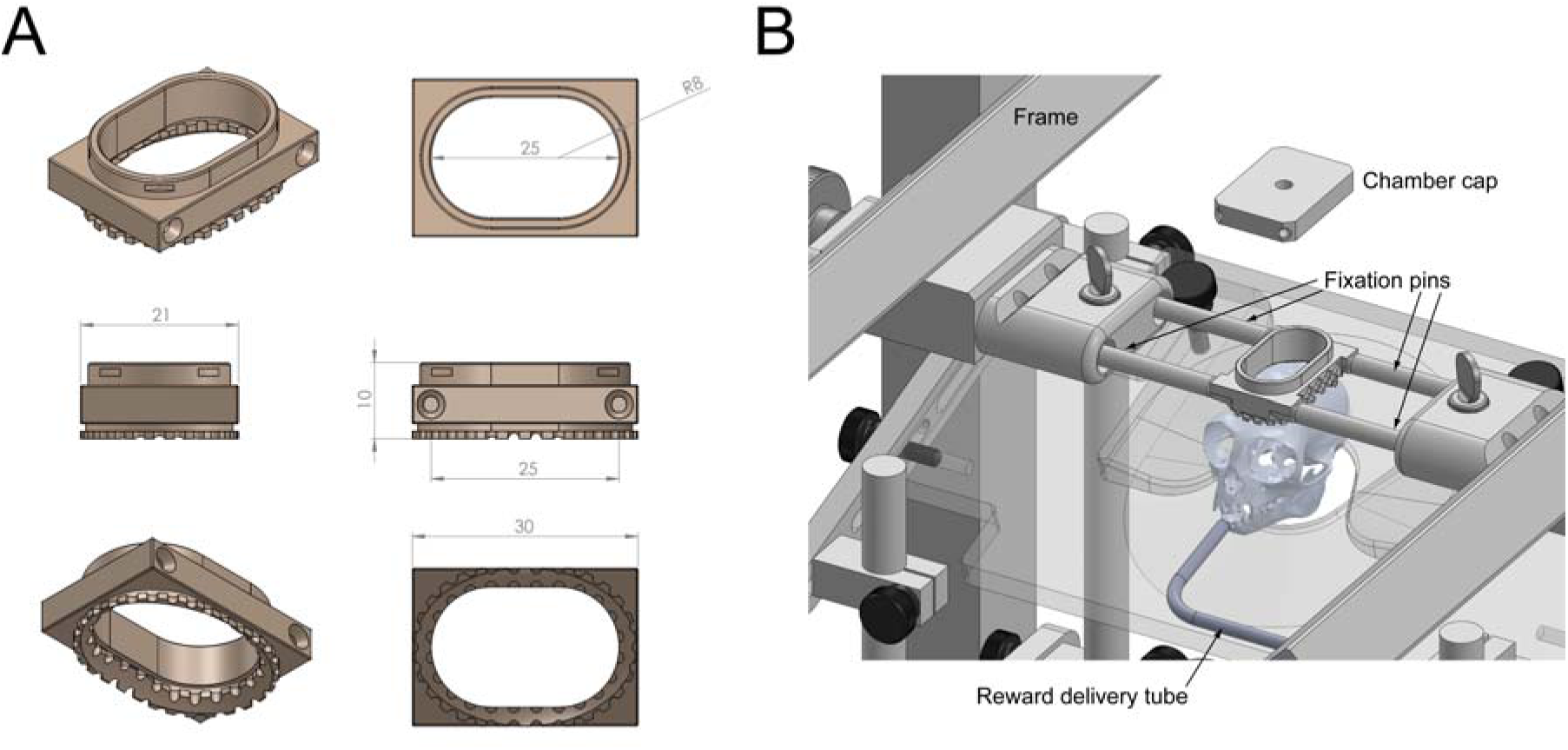
Custom fabricated combination head restraint/recording chamber. A) Schematics of chamber design with dimensions shown (mm). B) Schematic depiction of marmoset skull *in situ* within restraint chair, with head restraint/recording chamber, and head fixation pins shown. Metal tube in front of mouth allows delivery of liquid reward.

### Restraint Chair Design

Most marmoset restraint chairs used for neurophysiological and behavioural neuroscience studies are roughly based upon the early design of Hearn (1977), and use a common plan consisting of a tube surrounding the marmoset’s body for restraint, a baseplate to support the animal’s legs, and an adjustable plate which fits around the neck just above the shoulders to prevent the animal from escaping. For neurophysiological studies, a head holder is also used (O’Byrne & Morris, 1988; Wang et al, 1995; Remington et al., 2012; Mitchell et al., 2014). Our design differed in that we did not use a tube, but rather an adjustable waist plate in combination with an adjustable neck plate to restrain and position the animal’s body within the chair. The restraint chair we used here is depicted in Figure 2. We used this approach to allow greater freedom of movement of the marmoset’s arms, thus allowing the possibility of combining a touchscreen with head restraint in future studies combining touchscreen tasks and recording with moveable electrodes. Removable plastic shields can be inserted into the chair both above and below the waist plate to confine movements of the arms and legs within the chair for oculomotor studies. Both the waist and neck plates (.stl files are available for download), as well as a base grill to support the legs, are adjustable in relative height and angle via thumbscrews to provide a customized and comfortable fit for individual animals. This is desirable as minimizing discomfort is ethically sound practice and ensures a minimum of body movements, which can create unwanted artifacts during recordings. A stainless steel reward tube can also be attached to the chair and fed via gravity fed or injection pump to deliver liquid rewards that the animal can collect by licking.

**Figure 2.**
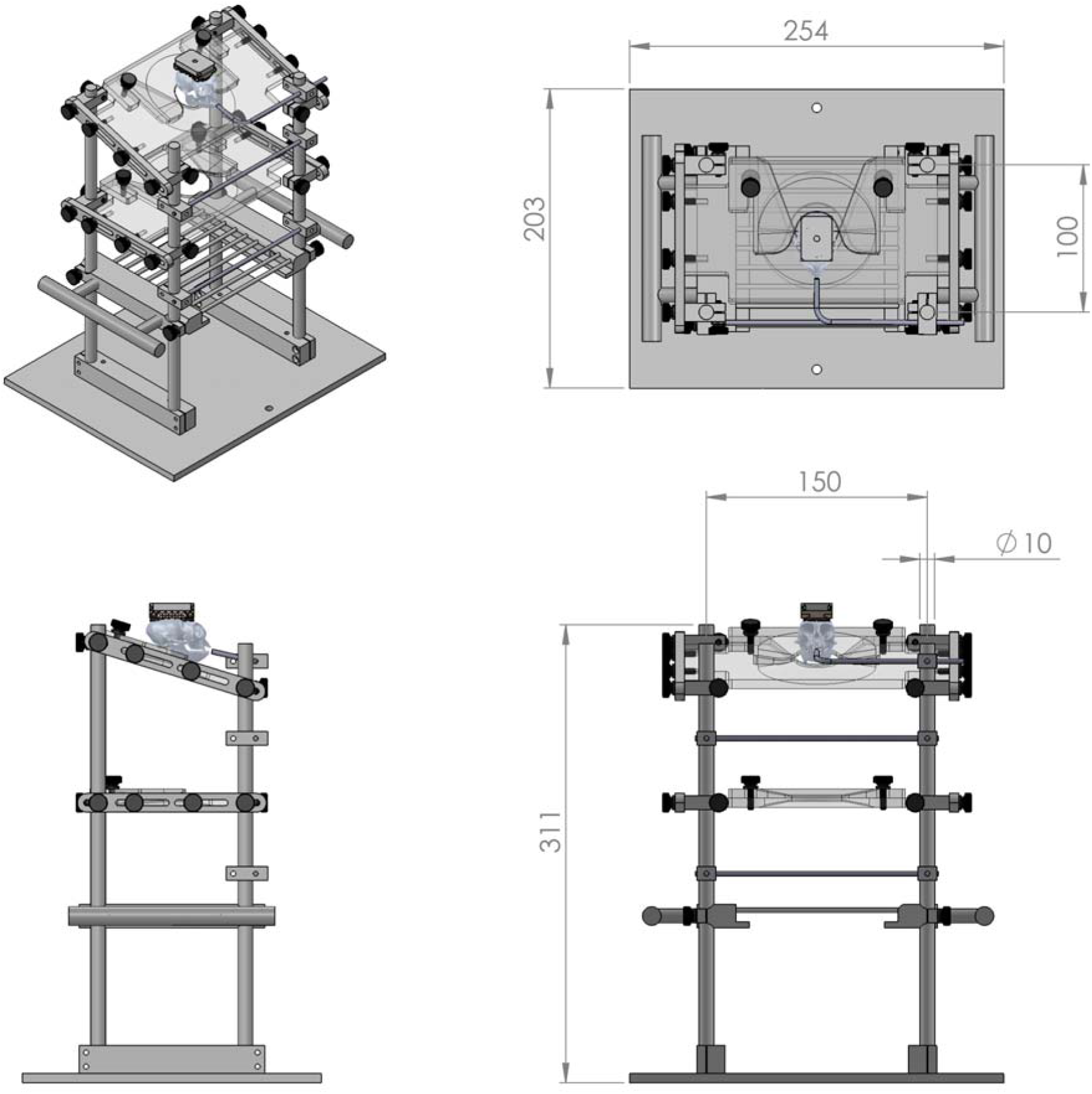
Schematic depiction of marmoset restraint chair, with dimensions (mm) shown in multiple views. Marmoset skull shown *in situ* as in Figure 1, for relative scale.

This chair system is designed to integrate with a custom designed stereotaxic frame for head restraint and mounting of microelectrode drives. The chair containing the marmoset can be easily slid through the back of the frame to position the animal, and clamped in place. This system is depicted in Figure 3. Altogether, this combined unit provides stable, comfortable restraint of the body and head, and a platform for mounting microelectrode drives or microinjection units for studies investigating pharmacological deactivation of brain areas.

**Figure 3.**
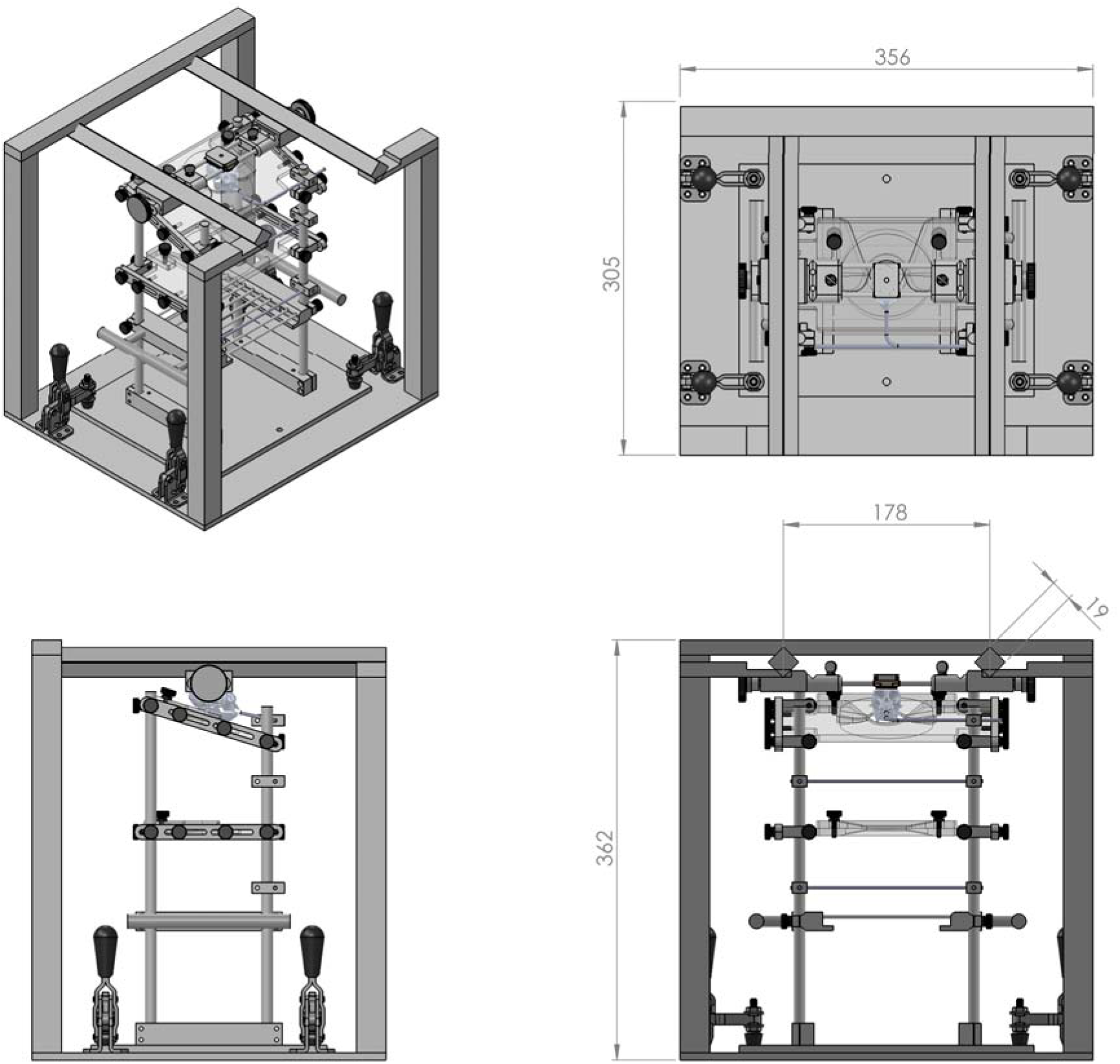
Schematic of marmoset chair within frame with dimensions (mm) in multiple views. Marmoset skull in position as shown in Figures 1 and 2.

In our lab, the chair/frame system is mounted on a table within a sound-attenuating chamber (Crist Instruments). A display monitor at the front of the chamber allows presentation of visual stimuli, and a high-speed video camera and infrared emitter (EyeLink 1000, SR Research, Ottawa, ON, Canada) are mounted below the monitor to enable video-oculographic eye movement recordings.

### Acclimation to Chair Restraint

Prior to implantation of the combination head restraint / recording chamber, we acclimated the animals to chair restraint. Prior to beginning any training on oculomotor tasks, it is imperative that the animal be trained to sit quietly and receive rewards.

We followed a procedure similar to that described by Silva et al. (2011) for fMRI studies in awake marmosets. Marmosets are brought to the experimental room in a transfer box, and hand transferred into the restraint chair. Within each session, the animals are presented with highly palatable reward items such as pudding, corn syrup, or acacia gum for remaining still and quiet. Initially, rewards are given for calm sitting in any position. Over sessions they are gradually shaped to sit facing forward by moving the presentation location of the food reward toward the front of the chair. Rewards are initially presented on a thin popsicle stick, and in the final stages, placed directly on the reward sipper tube at the front of the chair. In early sessions, the doors of the sound-attenuating chamber are left open, and the experimenter hand-delivers reward to the animal. In general the animals remain more calm if the experimenter remains in sight during the early stages of training. Once animals are able to consistently lick rewards from the sipper tube, the doors of the chamber are closed, and liquid rewards consisting of sweetened condensed milk, diluted corn syrup, or banana milkshake (selected according to the individual preference of each animal) are delivered via remote control of the reward pump while the experimenter monitors the animal via video camera.

The initial acclimation session is 10 minutes in length, and the duration of restraint is increased gradually in increments according to the progress of each animal until a total duration of 45 minutes is reached. Sessions are generally stopped at the first signs of the animal becoming agitated in order to avoid inducing undue stress. In our experience the amount of time required to complete this initial phase of training varies from between 6 to 8 weeks.

### Surgical procedures

Following acclimation to chair restraint, marmosets underwent an aseptic surgical procedure to implant the head restraint / recording chamber while the animal was placed in a stereotaxic frame (Kopf Instruments, Model 1248).

Anesthesia was induced with an intramuscular bolus of ketamine (15-20 mg/kg). A catheter was placed in one of the lateral tail veins and a continuous rate infusion (CRI) of propofol (0.3-5 mg/kg) in 0.9% saline was commenced, administered via a syringe pump. General anesthesia was additionally maintained with gaseous isoflurane (0.5-3.0 %) in oxygen, delivered through a custom-designed 3-D printed mask attached to the palate bar of the stereotaxic frame (see Supplementary Materials). Glycopyrolate (0.005-0.01 mg/kg) was administered to reduce salivary and bronchial secretions, and a loading dose of the NSAID meloxicam (0.2 mg/kg) given to reduce inflammation. An antibiotic, chosen based on the results of a pre-surgical swab and culture from each animal was also administered. A small amount of sugar paste was placed under the tongue to prevent hypoglycemia during surgery. Temperature was maintained with a circulating water blanket, forced-air warming blanket, and IV fluid warmer. Heart rate, SpO_2_ (measured by pulse oximetry), and temperature were monitored throughout the surgical procedure. A stereotaxic device was used to stabilize the head for placement of the combination head holder / recording chamber. A midline incision was made along the cranium and the temporalis muscle retracted to gain access to the underlying periosteum, which was carefully removed along with any tissue, and the surface was cleaned with (2%) hydrogen peroxide and cotton swabs. The skull surface was then prepared for application of dental crown cement. The “halo” type design we have employed allows access to a large area of the skull inside the chamber, which is beneficial for recording but occupies most of the lateral extent of the skull, making implantation of bone screws to secure the implant impractical due to space limitations. We have therefore used, with success, procedures similar to those applied in securing dental restorations, which rely on securing devices based on adhesive strength alone. This requires substantial preparation of the skull for optimal adhesion. We first stopped any bleeding on the skull surface using 15.5 *%* ferric sulfate hemostatic gel. The surface was mechanically abraded etched using a small metal brush (Dremel). Following this, the surface was thoroughly rinsed and any fluids or remaining bone residue were removed using absorbent swabs. Several coats of adhesive resin (BISCO ALL-BOND Universal, Schaumburg, Illinois, USA) were then applied using a microbrush, air dried, and cured with an ultraviolet dental curing light (King Dental). Adhesive cement (BISCO Duo-link, Schaumburg, Illinois, USA) was then applied to the skull surface and the head holder / recording chamber was lowered onto the surface using a stereotaxic manipulator to ensure correct location and orientation. Additional adhesive was added as needed to cover the skull surface inside the chamber and ensure an adequate seal around the chamber edges. Particular care was taken in smoothing the adhesive around the outside edges of the chamber to prevent any irritation of surrounding tissues. The adhesive was then cured with the dental light. If necessary, the rostral and caudal ends of the incision were closed with wound clips (Fine Science Tools) to ensure a tight interface between the wound edge and cement surface.

The animals were then recovered, and administered buprenorphine (0.01 - 0.02 mg/kg) for post-surgical analgesia before being placed in a warm, padded recovery cage. Buprenorphine treatment was continued for 2-3 days post surgery, along with administration of meloxicam (0.2 mg/kg) for up to 5 days, and an antibiotic regimen, all of which were evaluated and directed by a University veterinarian.

Neural recordings require a separate surgery to drill burr holes allowing access to the cortical surface, which follows training on oculomotor tasks. In this surgery, 3mm burr holes were drilled through the adhesive cement and skull inside the head holder / recording chamber using a micro-drill press mounted on a stereotaxic micromanipulator (Kopf Instruments).

### Eye movement recording and Stimulus Display

Eye positions were monitored via infrared tracking of the pupil at a resolution of 1000 Hz (EyeLink 1000, SR Research, Ottawa, ON, Canada). This system was arranged such that the camera and infrared emitter were mounted directly below the CRT video display on which visual stimuli were presented, at a distance of 42 cm from the animal. Some previous studies have noted that video tracking of the marmoset pupil can be difficult due to the large size of the pupil relative to the orbit. To compensate for this, visual stimuli have been presented on high luminance backgrounds, to ensure that the adaptation state of the animals is such that the pupil remains small enough to be tracked accurately (Mitchell et al., 2014), We did not find this to be the case with our system, and were able to obtain accurate tracking with the low background luminances we used here.

All stimulus generation and behavioural paradigms were under control of the CORTEX real-time operating system (NIMH, Bethesda, MD) running on Pentium III PC’s. Visual displays were presented on a 21” CRT monitor (ViewSonic Optiquest Q115, 76 Hz non-interlaced, 1600 x 1280 resolution). Stimuli consisted of a small circular fixation spot (0.25°, luminance 10 cd/m^2^) and larger circular target (0.8°, luminance 10 cd/m^2^), presented on a dark background (2 cd/m^2^).

### Training marmosets to perform oculomotor tasks

Marmosets were trained to make goal-directed saccades using methods adapted from those commonly employed for similar studies in macaque monkeys. We employed a method of successive approximations in which oculomotor behaviour progressively closer to that desired was rewarded. This proceeded in two stages: *initial fixation training,* in which the animals were trained to acquire and maintain gaze fixation upon suddenly appearing stimuli, and *saccade training,* in which they were trained so maintain gaze upon a fixation stimulus, and subsequently saccade to the location of a suddenly appearing visual stimulus. These phases will be detailed below. We noted that in these animals, it is imperative that a highly palatable reward is chosen to ensure consistent behaviour. We evaluated this in separate sessions by presenting the animals with varying liquid rewards and noting appetitive behaviours, such as licking frequency, for each. Rewards tested included acacia gum, corn syrup, maple syrup, banana milkshake, and sweetened condensed milk. We found that the most palatable reward differed between the two animals, with Marmoset B preferring sweetened condensed milk mixed with plain water in a 2:1 ratio, and Marmoset M preferring corn syrup diluted with plain water to a 1:1 ratio. We did not employ a fluid deprivation schedule in any of our training or data collection, but did use moderate food restriction. We reduced the amount of the animals’ second of two daily meals to 80% of their ad libitum consumption, and carried out all training or data collection sessions in the morning of the following day, prior to feeding of the first of two daily meals. Training sessions were conducted daily on weekdays, and no restriction was imposed on weekends. Both animals maintained body condition on this schedule.

#### Fixation training

For this phase of training, we exploited the previously reported preference of marmosets to shift gaze toward face stimuli (Nummela et al., 2017; Mitchell et al., 2014). An image consisting of a marmoset face, subtending 0.8 × 0.8° of visual angle, was presented at the centre of the display monitor, and the animals were rewarded for shifting gaze to, and maintaining gaze within a 5 × 5° electronic window centered upon this stimulus within 4000 ms to receive a liquid reward. The fixation duration requirement was gradually increased from an initial duration of 20ms to 500ms, in increments based upon the animal’s success rate and chosen by the experimenter. Trials on which the animals failed to acquire or maintain fixation were followed by a 5000ms “time out” period. Typically, steady fixation for the full 500ms duration was achieved within three to five daily sessions. Once performance was accurate with the fixation stimulus at a single, central location, we included a second location, 5 degrees to the left of fixation, with same spatial and duration requirements. Fixation trials at this location were randomly interleaved with the central location. Once the animals were able to reliably fixate at these two locations, a third location, 5 degrees to the right of fixation was added. Spatial and duration requirements were maintained as before, and trials at this location were interleaved with the central and left locations. Once the animals were able to fixate at all three locations for 500ms, we reduced the size of the electronic window surrounding the stimulus to 3 × 3°, and proceeded to saccade training.

#### Saccade training

Once animals were trained to fixate three horizontal stimulus locations for 500ms within a 3 × 3° window, we proceeded to saccade training. This phase of training used the same face stimuli as in fixation training. The animals were presented with a central fixation stimulus, upon which they were required to maintain gaze for a duration of 500ms. If fixation was successful, the central stimulus was extinguished, and replaced immediately by a second stimulus presented at 5° to the left of fixation. If the animal shifted gaze to the new location within 1000ms and maintained fixation for a duration of 10ms within a 5 × 5° window surrounding this stimulus, a liquid reward was delivered. In contrast to macaques, which readily shift gaze in response to such to such target “jumps”, we noted that these marmosets often maintained gaze at the central location despite the fact that no stimulus was present there, and that the peripheral stimulus was presented at an eccentricity well within the oculomotor range of the animals, which extends to 10 degrees (Mitchell et al., 2014). For this reason, and to encourage saccade behaviour, we employed the relatively generous criterion of 1000ms noted above. In the initial stages of training, we also limited to presentation of the peripheral stimulus to the left of fixation so that it’s location was predictable and to encourage saccadic responses. In both cases, the animals made saccades to the target location within 2-3 sessions. Following this, we added a second peripheral target 5° to the right of fixation, and ran each condition in 10 trial blocks. Once the animals were proficient at this, we randomized the stimulus locations and reduced the maximum duration allowed between fixation stimulus disappearance and saccade onset to 500ms. Following this, we replaced the face stimuli with dots as used in the step and gap paradigms. This stimulus generalization was not difficult for the marmosets and was completed in a single session.

### Behavioural paradigm

Marmosets performed saccades to peripheral targets in “step” and “gap” conditions. Schematics of these paradigms and representative eye traces are presented in figure 4. All trials began with the appearance of a small fixation spot at the centre of the screen. Animals were required to look at this stimulus within 3000 ms of its appearance, and to maintain gaze within an electronic window of 1.5° x 1.5° surrounding it for 700 – 900 ms. On “step” trials a peripheral target stimulus, was presented on the same horizontal meridian as the fixation stimulus, pseudorandomly to the left or right of fixation, coincident with offset of the fixation spot. Animals were required to generate a saccade toward the peripheral stimulus within 500ms of it’s onset, and were given a liquid reward if the saccade endpoint fell within a 3° × 3° electronic window centered on the target location (Fig. 4A). On “gap” trials, animals were required to maintain fixation on the fixation spot for 500 – 700ms. Following this, the fixation spot was extinguished for a “gap period” of 200ms, during which the animals were required to maintain gaze within the electronic window previously centered on the location of the fixation spot. At the end of the gap period, a peripheral target stimulus was presented pseudorandomly to the left or right of fixation, and the animal was given liquid reward for meeting the same behavioural requirements as the “step” condition (Fig. 4B). Following correct trials, the intertrial interval lasted 2000 ms.

### Data Analysis

All analyses were carried out in MATLAB (Mathworks, Natick, MA) using custom-written software. Saccade onset and offset were defined at the times at which the horizontal eye velocity exceeded or fell below 30°/sec, respectively. Trials with saccadic reaction times (SRTs) below 50ms were classified as anticipatory rather than stimulus-driven saccades. Those with SRTs exceeding 500ms were classified as no-response trials. Both of these trial types, as well as those with incorrect or broken fixation, were excluded from further analysis.

## RESULTS

### Combination head restraint/recording chamber

Both animals implanted using the methods described here have maintained their implants, Marmoset M for 16 months, and Marmoset B for a duration of 8 months. During this time, routine maintenance of margins surrounding the outside edges of the chamber, including trimming of hair and cleaning of the skin with hydrogen peroxide, have been sufficient to maintain healthy tissue without infection. In neither case did we observe any irritation or reaction of the skin to the dental cement used here. Marmoset M was subsequently a participant in a second study in which chronic neural recordings were carried out in both frontal and parietal cortex during performance of saccade tasks. In this case 16 contact linear electrodes (Atlas Neuroengineering, Leuven, Belgium) were driven into the brain using stereotaxic arms attached to the frame used for head restraint. These data will be presented in detail in a subsequent report, however Figure 5 presents a schematic of the electrode and local field potentials aligned on visual stimulus onset from a single trial of recordings in posterior parietal cortex to provide proof of principle. We found that the chair and head restraint system described here provided a stable platform for neurophysiological recordings during oculomotor tasks.

**Figure 4.**
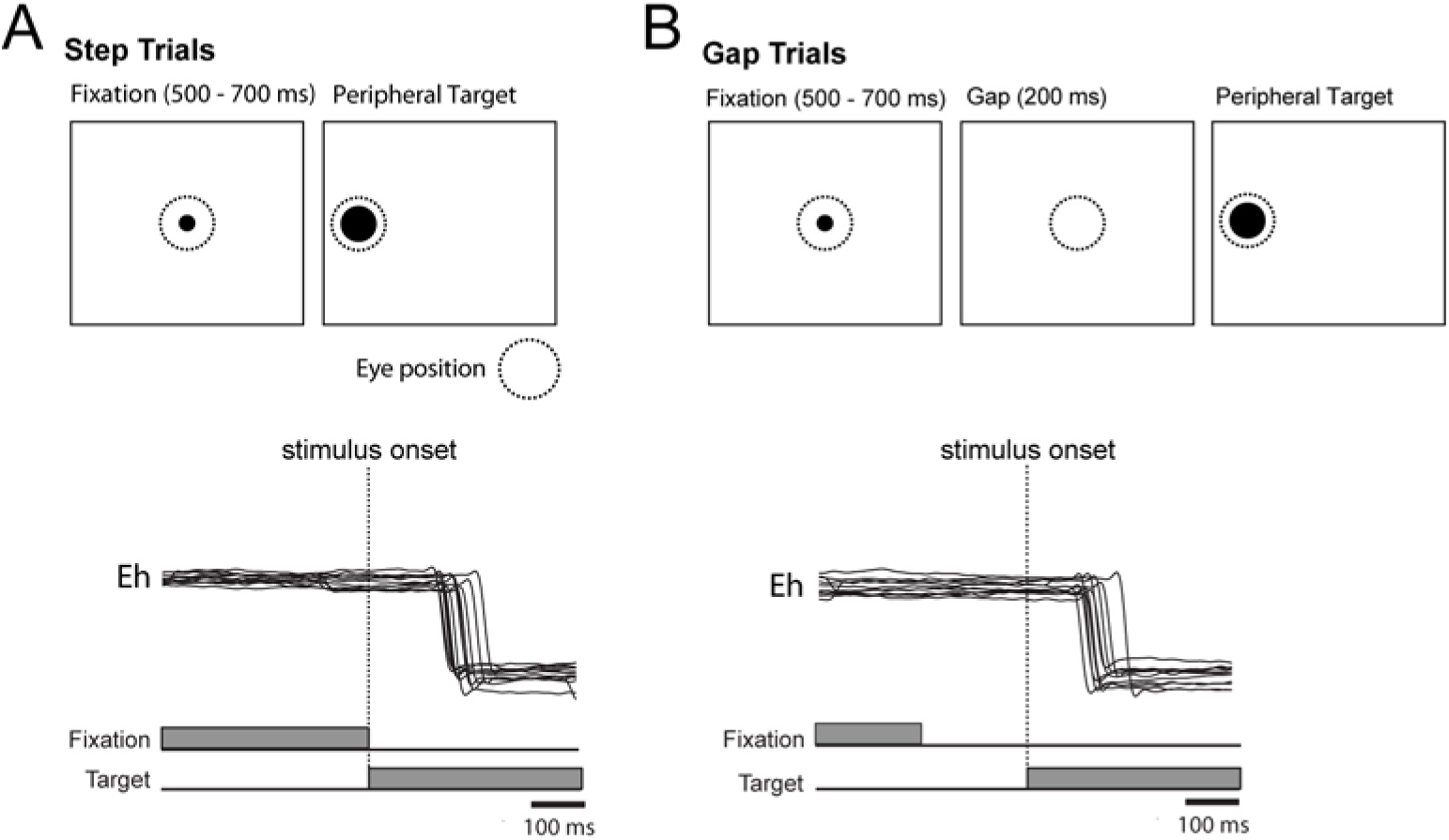
Schematic depiction of oculomotor tasks. A, upper panel, step trials. After initial fixation period, marmosets were rewarded for generating a saccade toward a peripheral target stimulus which appeared coincident with offset of the fixation stimulus. Lower panel, timeline of fixation and target stimulus appearance, and representative eye traces. B, upper panel, gap trials. Marmosets were rewarded for generating a saccade toward a peripheral target stimulus following initial fixation and a 200ms “gap” interval during which the fixation stimulus was absent. Lower panel, timeline of fixation and target stimulus presentation and representative eye traces.

**Figure 5.**
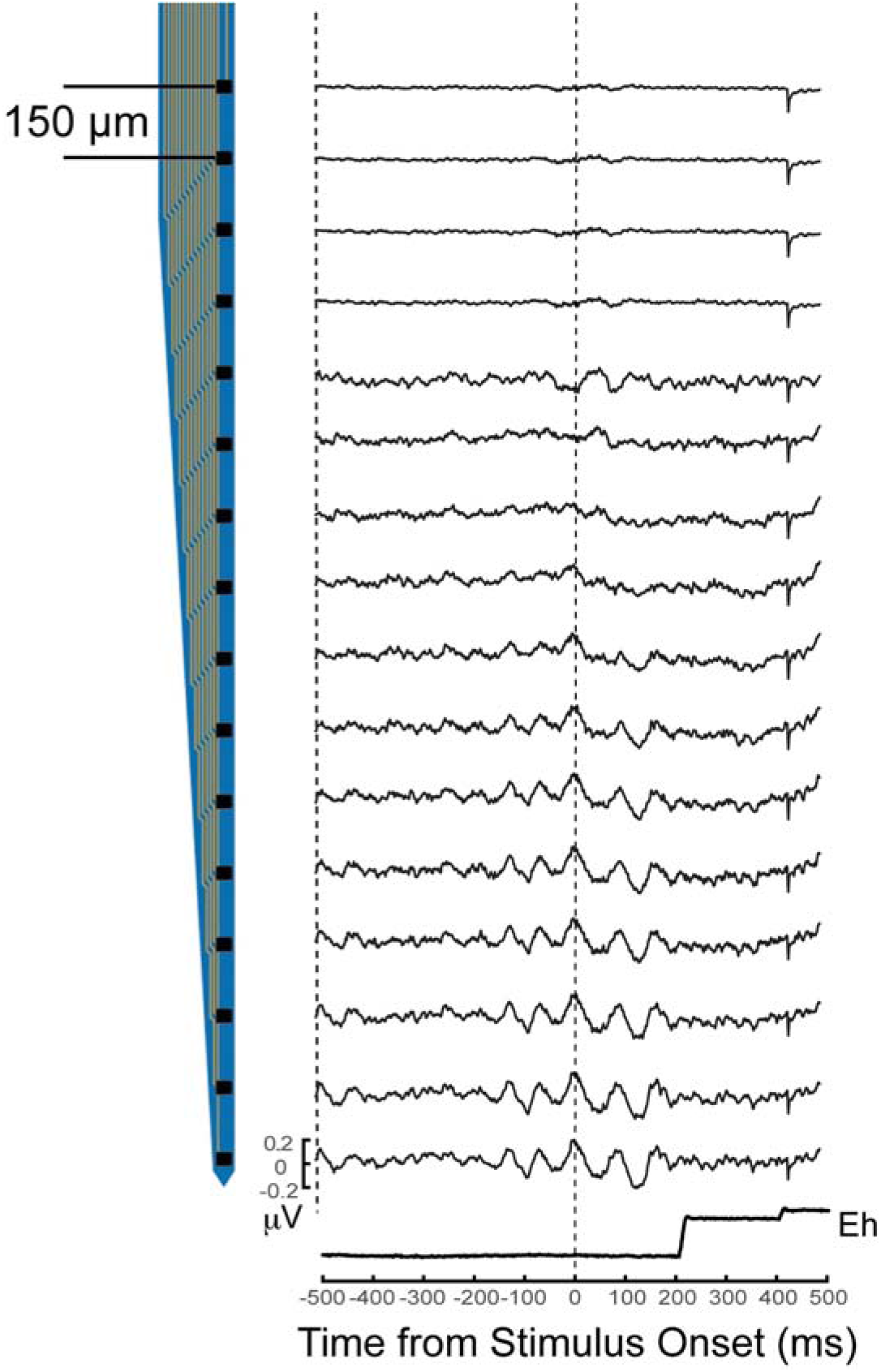
Schematic representation of linear recording electrode and trial local field potentials (LFPs) recorded from posterior parietal cortex of Marmoset B. Left, schematic of electrode depicting each of 16 contacts. Right, LFPs recorded on each contact during a single trial in which marmoset made a rightward saccade. Data are aligned on onset of peripheral visual stimulus, timeline shown with eye trace at bottom right. Scale bar on LFP signals denotes +- 0.2 μV.

### Training outcomes

Using the methods described here, we were able to successfully train both marmosets to perform the step and gap saccade paradigms over a time period of approximately 5 months. Of this, the initial acclimation to the restraint chair accounted for 6-8 weeks of the training period, and the remainder for training the animals to maintain stable fixation (2-3 weeks), generate goal-directed saccades (4 weeks), and perform the step and gap tasks (4 weeks). We present data from both animals in these paradigms below. During data collection, we found that animals reached satiation after a period of approximately 25-35 minutes in each session.

### Performance of Step and Gap Tasks

Both Marmoset B and Marmoset M were proficient at these tasks. Data presented for Marmoset B were collected over 4 sessions with a total of 381 trials. Data for Marmoset M were collected over 6 sessions in which the animal performed a total of 542 trials. Of these, 349/381 and 528/542 remained for analysis after applying our exclusion criteria. We observed a significant reduction in reaction times on gap as compared to step trials for both animals. SRT histograms for step and gap trials for Marmoset B (panels A, B, respectively) and Marmoset M (panels C, D, respectively) are presented in Figure 6. We found that the SRT distributions for both trial types in both animals were significantly non-normal (Kolmogorov-Smirnov tests, p < .005 in all cases), and thus carried out statistical comparisons of median SRTs using nonparametric Wilcoxon ranksum tests. For Marmoset B, we observed a statistically significant reduction in median SRTs between step and gap trials (M = 108 ms vs. M = 98 ms, Z = 3.21, p = .00027). We observed a similar but larger reduction in median SRTs between step and gap trials for Monkey M (M = 162 ms vs. M = 129 ms, Z = 8.90, p = .00083. Taken together, these data provide evidence for a gap effect in marmoset SRTs, and demonstrate that this primate species can be trained to perform at least basic oculomotor tasks using the methods described here.

**Figure 6.**
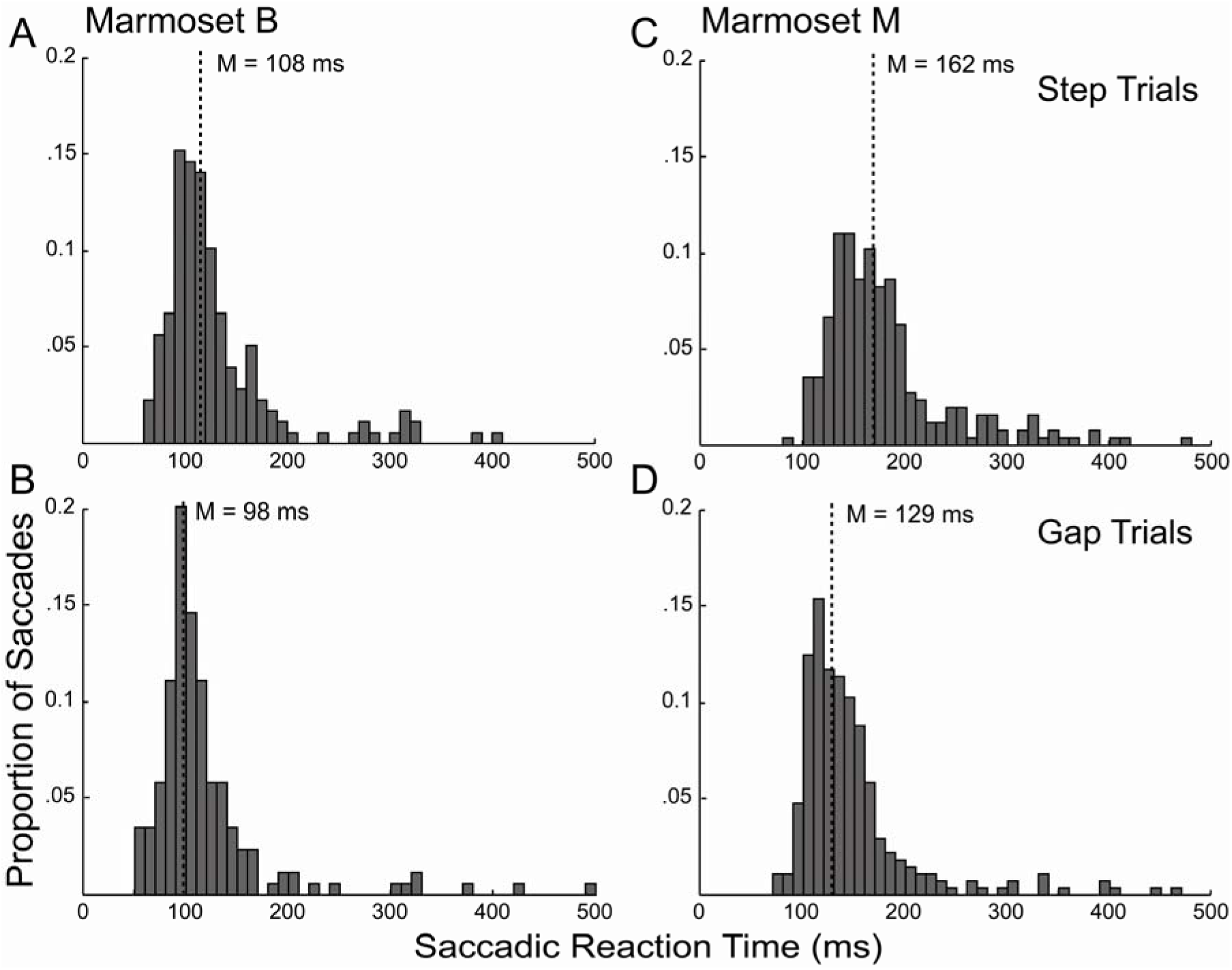
Saccadic reaction time (SRT) histograms for Marmosets B and M on step and gap trials. Panels A and B depict SRT’s for Marmoset B on step and gap trials, respectively. C and D, same as A and B, but for Marmoset M.

## DISCUSSION

Neuroscience research using the common marmoset model is expanding rapidly due to the development of methods allowing unique insights into the operation of neural circuits in a primate brain in health and disease. The ongoing adaptation of techniques developed in rodents to research in primate brains, such as two-photon calcium imaging of small populations of individual neurons (Sadakane et al., 2015), optogenetic approaches to manipulate neural activity (MacDougall et al., 2016), and the possibility of developing transgenic models of brain disorders (Jennings et al., 2016), promise significant advances in our understanding of neural circuits and their modifications in disease and health. In parallel, methods for training and carrying out neural recordings in behaving marmosets are advancing. Neural recordings in awake behaving macaque monkeys have contributed immensely to our understanding of sensory, motor systems, and cognitive systems. Combining the power of this technique with those available in the marmoset is an exciting prospect. Here, we have detailed our adaptation of surgical methods for implantation of a novel, custom designed head restraint/recording chamber, design of a custom marmoset chair and methods of acclimation to chair restraint, and methods of training on saccade tasks to the marmoset model, and successfully used these methods to train and collect data from two marmosets during performance of a visually-guided saccade task.

Previous work in this species has used differing methods of head restraint and of obtaining access to the brain for neural recordings. For example, Lu et al. (2001) used a head post for head fixation, and small burr holes drilled in the skull and surrounded by a well of dental adhesive for neural recordings. Similarly, Mitchell and colleagues (Mitchell et al., 2015; Mitchell et al, 2015) have also used a head post system. This system requires that an adequate amount of adhesive and bone screws are used to ensure that this post remains stable, which in turn requires a significant amount of space on a relatively small skull. As our primary interest is in simultaneous recording from multiple cortical areas, this lack of available recording space presents a potential limitation, and thus we used the “halo” design described here. We were able to achieve stable head restraint throughout training and data collection with this system, and this method allowed us to record through burr holes drilled in the skull over locations of interest. This method combines the possibility to access a larger area of cortex with the known stability of recording through small burr holes in this species (Lu et al., 2001).

In contrast to other studies using marmosets, we found that our animals performed fewer trials per session before reaching satiation. Our animals performed 80 -100 trials per session, as compared the 300 – 800 reported by Mitchell and colleagues (2014). Similar to these authors, we also employed a moderate food restriction regimen for one of our animals to control motivation, and thus we do not believe that this was a factor. We attribute this difference to the amount of reward delivered on each trial. For each animal, we adjusted the volume of reward delivered per trial at the beginning of training to maximize the animal’s work rate, and did not subsequently reduce the amount provided once they had learned the tasks. The amount of reward delivered on each trial was 0.07 ml, roughly 0.05 -0.06 ml more per trial than that given in the study by Mitchell et al. (2014). At this reward amount, each animal would satiate relatively quickly, and consume roughly 7 ml in a 20-30 minute session, an amount within the range of the 5-15ml per session reported in their study. To address this, we plan to reduce the per-trial reward amount for trained animals, and may replace the gravity-fed reward system with a precisely calibrated fluid mini-pump to optimize the amount of reward delivery. Notwithstanding this, we were able to collect a statistically significant dataset in only a few sessions for each animal.

We found that, similar to humans and macaques, a psychophysical gap effect on saccade latencies was exhibited in the marmoset. The values we obtained here, 10 ms for Marmoset B, and 33 ms for Marmoset M, were consistent with previous studies in macaques (11-48 ms, Krauzlis & Miles, 1996; 19.7 - 34.9 ms, Paré & Munoz, 1996), though somewhat shorter than those seen in studies of human participants, which average roughly 60 ms (Saslow, 1967; Fischer & Boch, 1983). Neural correlates of the gap effect have been identified in the discharge properties of single neurons in the intermediate layers of the SC. Specifically, it has been proposed that the characteristic reduction in reaction times results from the dynamic interplay of neurons in the rostral SC, related to gaze-holding, and those in the more caudal SC, related to gaze-shifting (see for review Munoz et al., 2000). Fixation neurons in the rostral SC have been shown to decrease their rate of discharge during the gap period, following the offset of a central fixation stimulus (Dorris & Munoz, 1995), while saccade-related neurons in the caudal SC exhibit an increase in low-frequency activity related to saccade preparation during this time (Dorris et al, 1998). It has been proposed that this simultaneous release from active fixation and increase in saccade preparation accounts for the reaction time decrease observed on gap trials. An alternative account holds that the absence of visual stimulation of rostral SC neurons following offset of the fixation stimulus on gap trials and corresponding increase in SC saccade-related neurons, rather than release of an active fixation process *per se,* accounts for the gap effect (Krauzlis et al., 2017). Notwithstanding the details of the specific processes instantiated by rostral SC neurons, it seems clear that the gap effect is a behavioural manifestation of interactions across the SC map. The presence of this effect in the marmoset, together with the similarity in the magnitude of this effect between rhesus and marmosets, suggests that homologous SC circuits underlie this effect in primates, and supports the marmoset as a model animal for neurophysiological investigations of oculomotor control.

Due to it’s many demonstrated advantages, recent work has sought to adapt existing methodological knowledge from the macaque monkey model to operant conditioning of tasks and simultaneous neural recordings in the marmoset. Head-restrained marmosets have been trained to perform both auditory (Osmanski & Wang, 2011; Remington et al., 2012; Osmanski et al., 2013), and eye movement (Mitchell et al., 2014; Mitchell et al., 2015) tasks. An extensive series of studies of single-neuron activity in the marmoset auditory cortex have been carried out (Lu et al, 2001; Barbour & Wang, 2003; Bendor & Wang, 2005), and studies combining these methods (Bendor et al., 2012) are becoming more common. In the context of previous work showing similarities in oculomotor control between marmosets and macaques (Mitchell et al., 2014), our work adds additional evidence for the viability of operant conditioning methods for training of oculomotor tasks, the applicability of chair and head restraint to stable neurophysiological recordings during investigations of oculomotor control in the marmoset, and further support for the homology of saccade networks between these primate species. Studies combining these methods with further high-density and laminar recordings promise to add significantly to our knowledge of neuronal circuit operations underlying control of eye movements, and to shed light on the general principles of cortical processing.

## ACKNOWLEDGEMENTS

The authors thank N. Hague, K. Faubert, and A. Kirley for exceptional animal care and surgical support.

## SUPPLEMENTARY MATERIALS

**Supplementary Figure 1.**
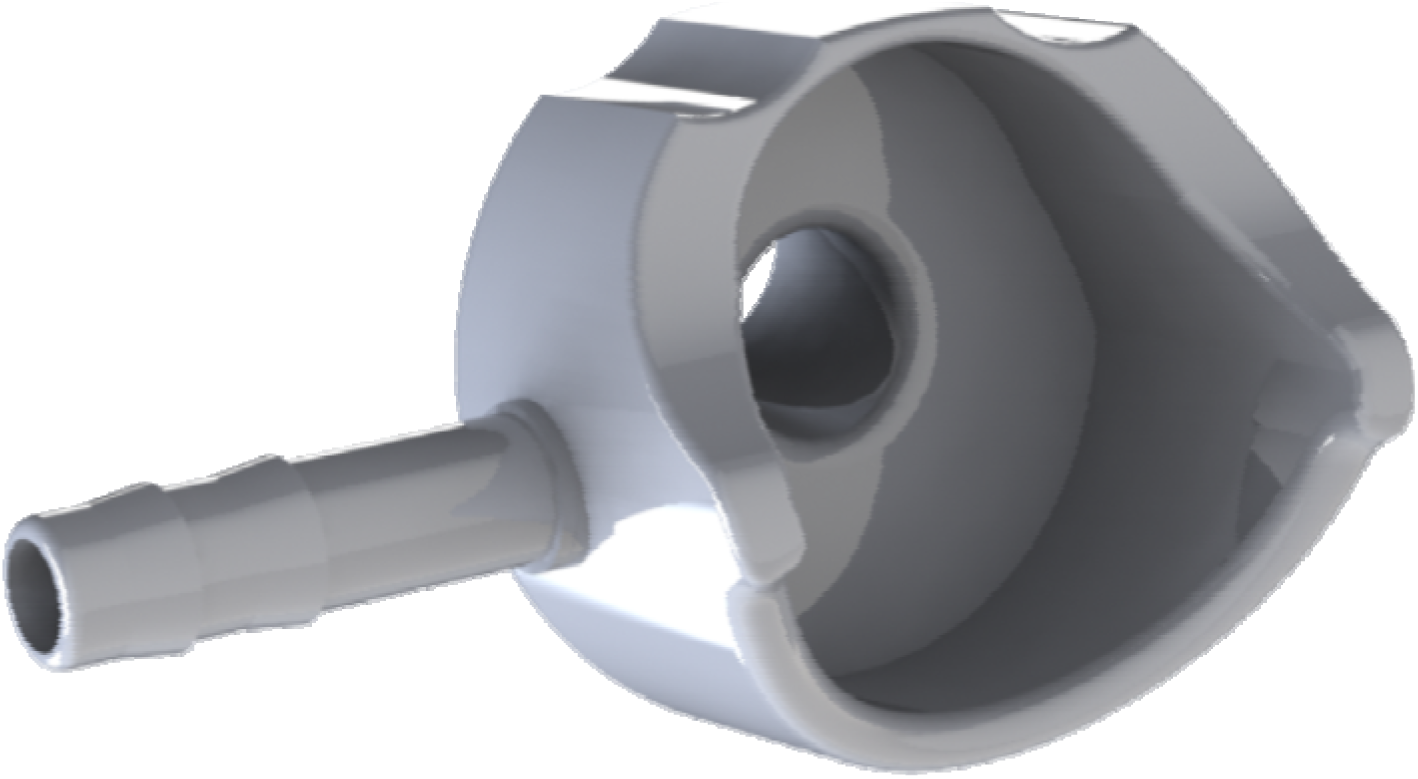
Custom-designed anesthesia mask for marmoset surgeries. Mask was designed using Solidworks (Waltham, MA) and 3D printed. When mounted on the palate bar of the stereotaxic frame, this allowed for a stable platform and precise fit, covering both the mouth and nares of marmosets, allowing controlled delivery of isoflurane and oxygen. An accompanying .stl file is available for download.

